## Supplementary figures and images for "Spatial and longitudinal tracking of enhancer-AAV vectors that target transgene expression to injured mouse myocardium"

### Figure S1

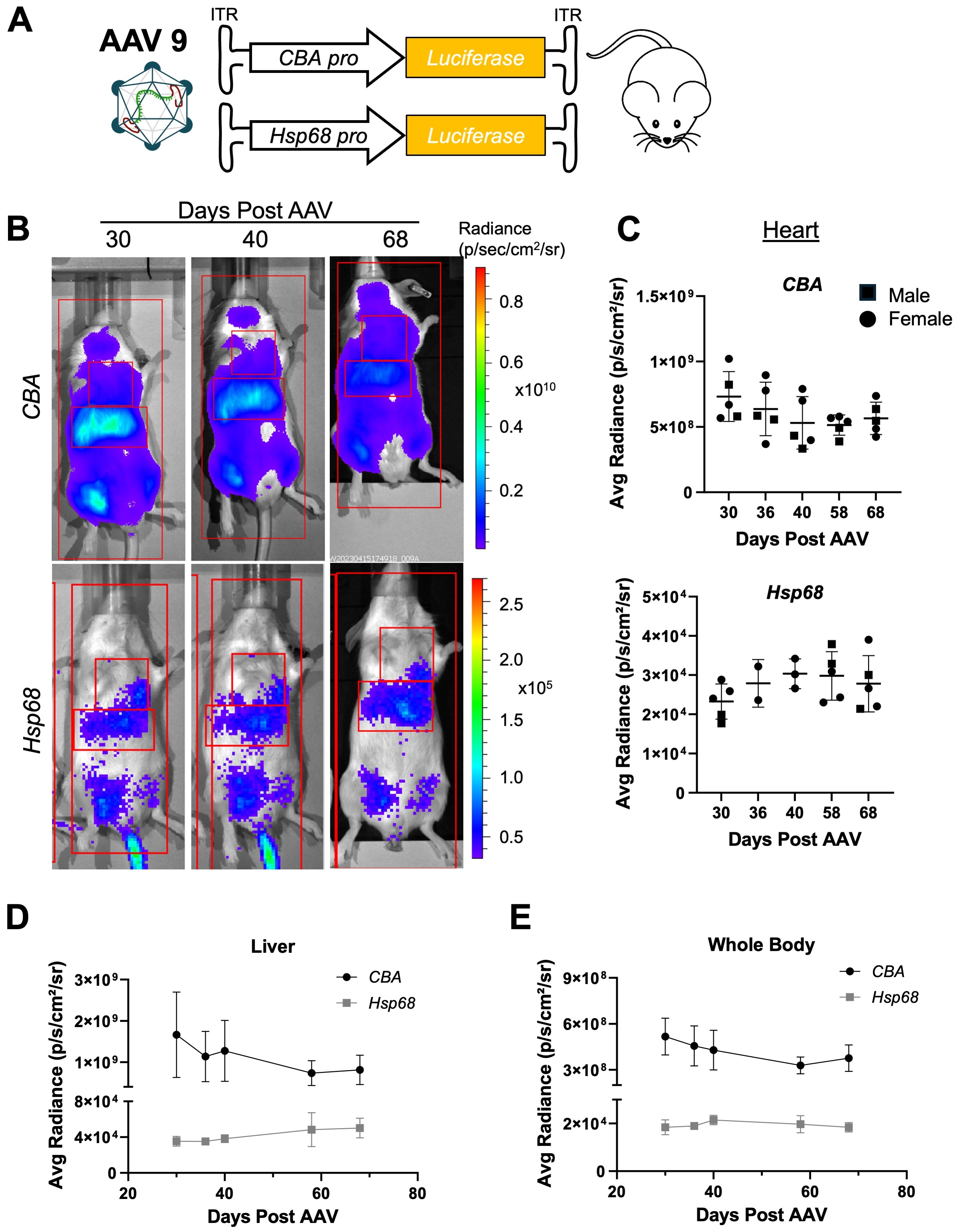

### Figure S2

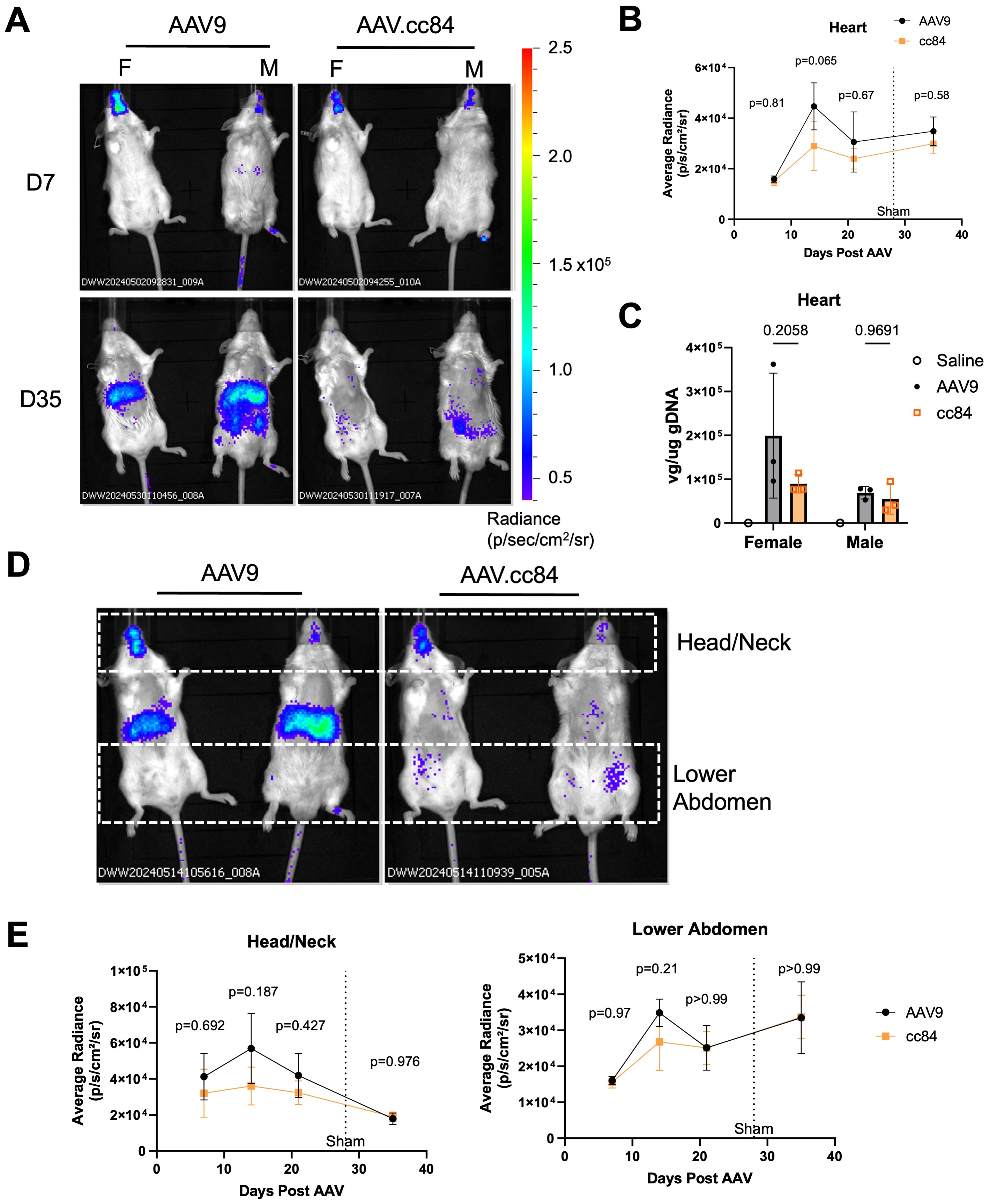

### Figure S3

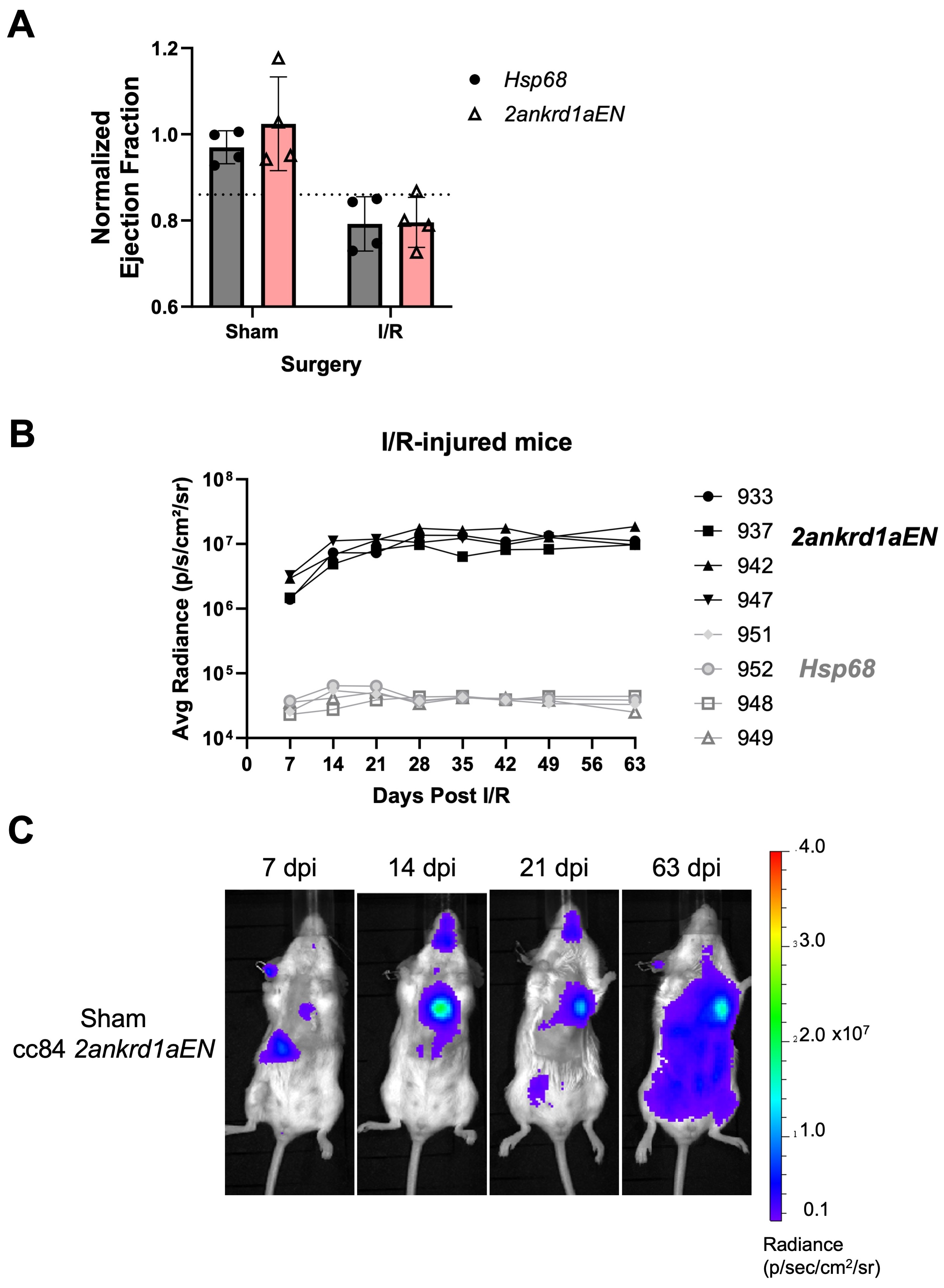

### Figure S4

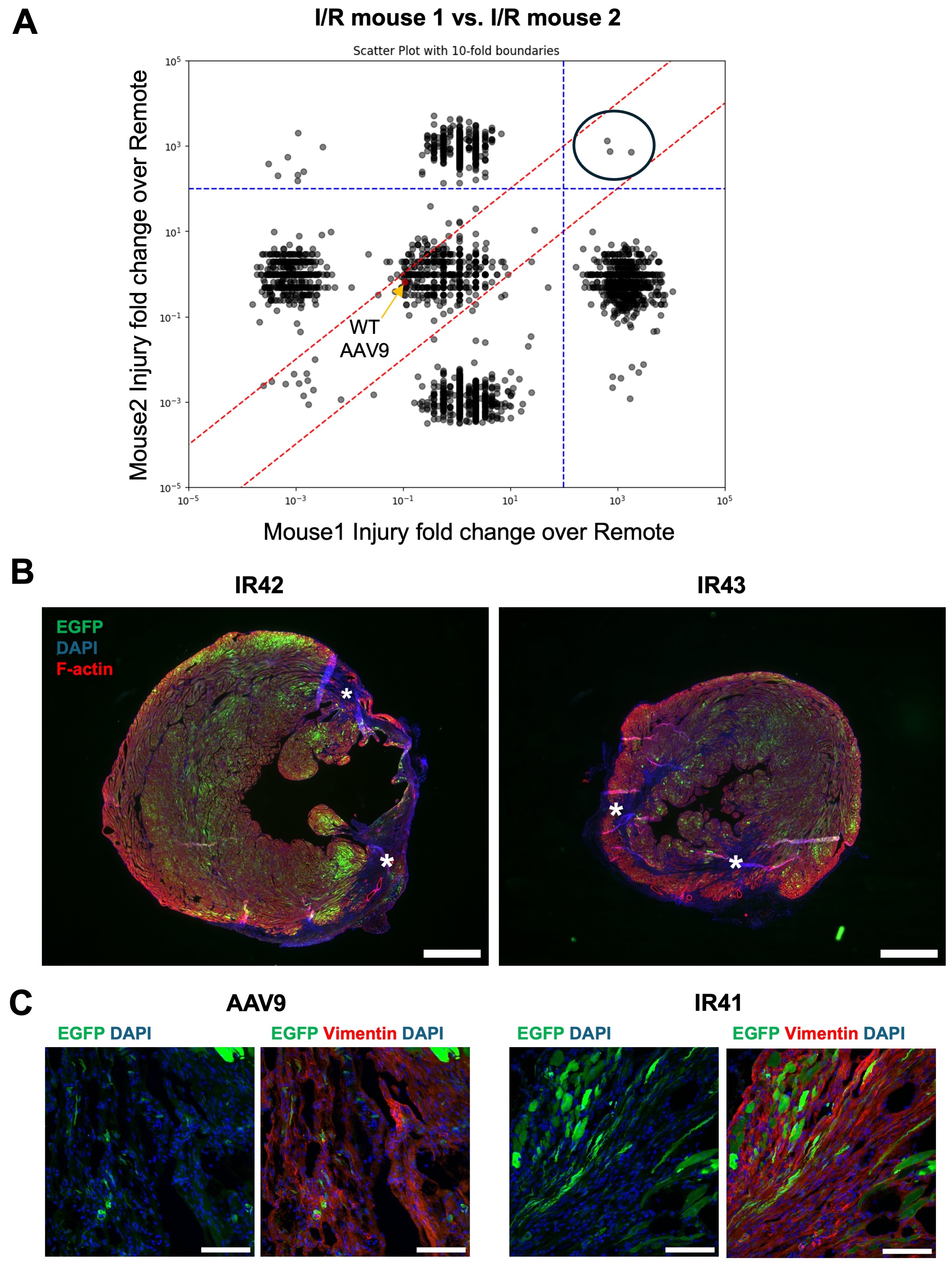

### Figure S5

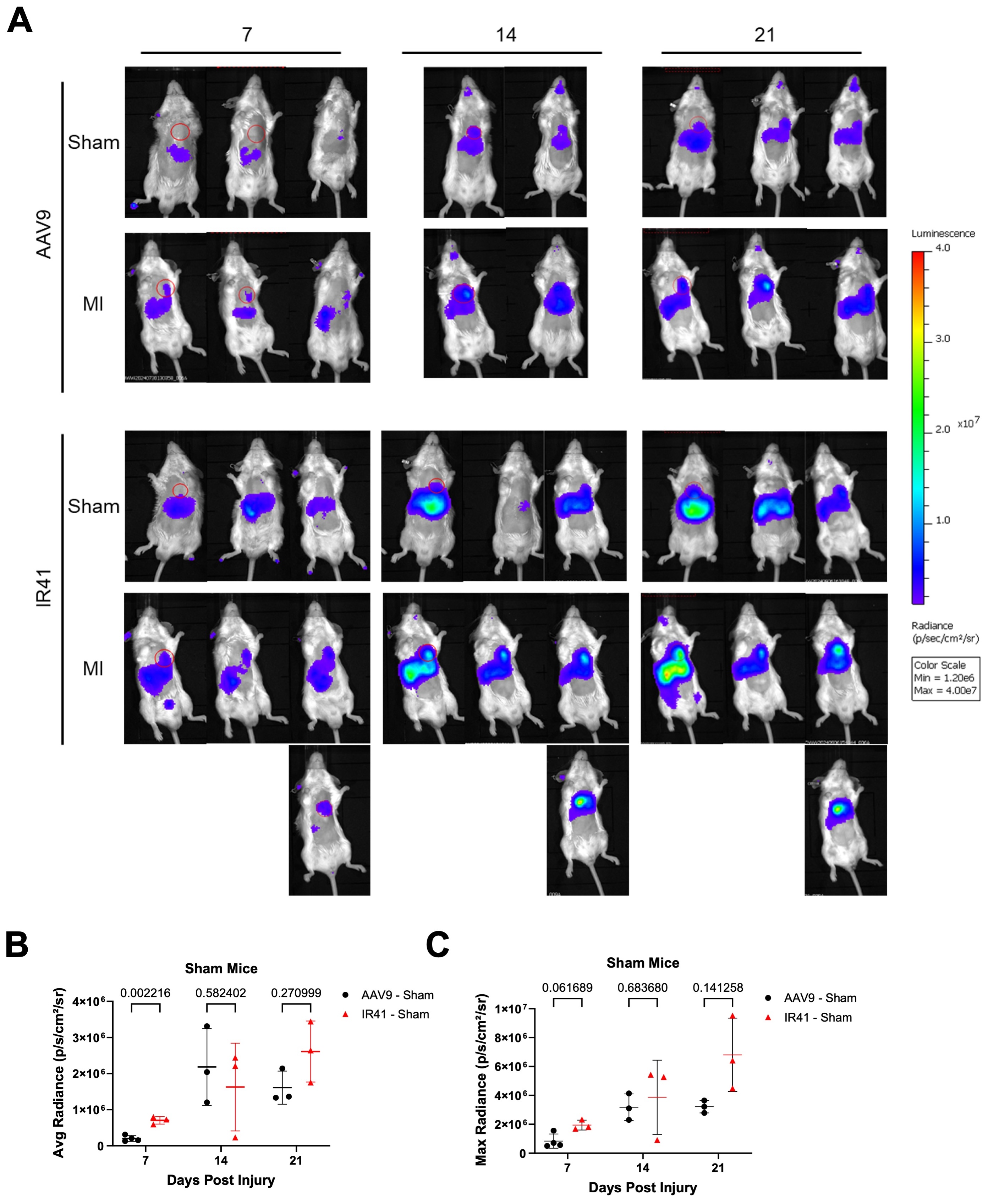
